# Intra-species diversity ensures the maintenance of functional microbial communities under changing environmental conditions

**DOI:** 10.1101/530022

**Authors:** Natalia García-García, Javier Tamames, Alexandra M. Linz, Carlos Pedrós-Alió, Fernando Puente-Sánchez

**Affiliations:** Systems Biology Program, Centro Nacional de Biotecnología, CSIC, C/Darwin n°3, Campus de Cantoblanco, 28049 Madrid, Spain; Great Lakes Bioenergy Research Center, University of Wisconsin-Madison, Madison, WI, USA

**Keywords:** intra-species diversity, microdiversity, ASV, ecotype, microbial community, environmental conditions, temperature, stability

## Abstract

Intra-species diversity comprises different ecotypes within the same species. These are assumed to provide stability in time and space to those species. However, the role that microdiversity plays in the stability of whole microbial communities remains underexplored. Understanding the drivers of microbial community stability is necessary to predict community response to future disturbances. Here, we analyzed 16S rRNA gene amplicons from eight different temperate bog lakes at OTU-97% and amplicon sequence variant (ASV) levels, and we found ecotypes within the same species with different distribution patterns in space and time. We observed that these ecotypes are adapted to different values of environmental factors such as water temperature and oxygen concentration. Our results showed that the existence of several ASVs within a species favored its persistence across changing environmental conditions. We propose that microdiversity aids the stability of microbial communities in the face of fluctuations in environmental factors.

## Introduction

Microbial communities have a profound effect on the planet’s surface and atmosphere. These communities are composed by hundreds to thousands of different species of microbes, but the nature of the relationships between microbial diversity and community and ecosystem functioning remains one of the big unanswered questions in general ecology [1]. Subtle relationships between the environment and microbial species can affect the importance of microbial community structure to explain ecosystem-level processes [2]. Moreover, high diversity has been tied to resilience in some microbial communities [3, 4].

Traditionally, species have been considered to be the units of diversity. In the case of microorganisms, the formal definition of a species requires isolation in pure culture, phenotypic characterization and sequencing of the genome [5]. Nevertheless, the concept and definition of microbial species are still open issues in microbiology leading to extensive debate [6, 7]. Besides, the usefulness of species as the most significant unit in microbial ecology is also controversial [8–10] due to the fact that two individuals classified within the same species may carry out different functions in the environment [11, 12]. Each species is formed by several strains that share a core genome and differ in the accessory genome. The term pan-genome has been coined to refer to all the genes that can be found in the known strains of a species, including the core and all the accessory genes [13]. Variations in the accessory genome are assumed to be responsible for niche differentiation and allow the discrimination of different ecotypes, pragmatically defined as populations of cells adapted to a given ecological niche [14–16]. When dealing with natural environments, it is not possible to sequence the complete genomes of all the populations involved, so proxies have to be used for species and ecotypes. The 16S rRNA gene can be used as a preliminary step to guide decisions regarding other steps involving less affordable sequencing technologies [17], and it has been used, in almost all cases, to define operational taxonomic units (OTUs). Commonly, OTUs have been defined at the 97% similarity level (assumed to correspond approximately to species), although an increasing number of authors considered this cutoff to be too low [18].

This perceived necessity for higher taxonomic resolutions was countered by the concern that, at very high similarity cutoffs, sequencing errors would lead to artifactual clusters [19]. Recent methodological developments, however, have overcome this limitation [20–25], allowing the discrimination of Amplicon Sequence Variants (ASVs) with only one nucleotide difference in their 16S rRNA gene sequence [26, 27]. ASVs make marker-gene sequencing more precise and reproducible than OTUs-97% [27]. Analyzing these ASVs may explain the distribution of ecotypes across environments, unveiling previously overlooked ecological patterns [20, 28–31].

The existence of ecotypes with overlapping ecological functions (but adapted to different niches) has been proposed to confer stability to microbial ecosystems [32], guaranteeing the long-lasting persistence of bacterial populations [33]. Indeed, several species have been shown to consist of many different subpopulations that sustain their distribution across broad environmental gradients [12, 30, 31, 34]. However, it is still not clear whether this intra-species diversity also contributes to the preservation of the higher order inter-species interactions that are fundamental for community functioning and stability. Stability has been studied extensively [35–37], especially in relation with diversity [36, 38]. Diverse communities will contain functionally equivalent organisms able to respond differentially to the environment, resulting in increased stability against environmental perturbations [36]. On the other hand, the stability of microbial communities is also influenced by the type and strength of the ecological interactions established among their members. Species capable of forming versatile interactions with multiple partners will be less affected by variations in community composition; a community dominated by such weak interactions will thus be more robust against change [36]. As microbial communities play a relevant role in ecosystem processes, understanding the drivers of microbial community stability is important to predict community response to future disturbances (for a thorough review see [39]), such as those resulting from global change.

Here, we have examined the identity and distribution of OTUs-97% and ASVs using a large dataset of samples collected in eight different temperate bog lakes in Wisconsin, U.S.A. [40, 41]. These lakes are all located in the Northern Highlands, a rocky, high elevation area in the southernmost edge of the North American boreal biome. Lakes have been studied intensively [42–46] because of the fact that these systems play an important role in human activities [47–49] and can be used as sentinels and integrators of environmental change [50]. Besides, the fact that lakes are relatively similar to each other, but with differences in environmental parameters such as temperature, oxygen, or mixing regimes, make these systems fitting to address some core questions in microbial ecology such as how community assembly takes place, how interactions among taxa influence community composition, or how microbial communities respond to environmental perturbations such as mixing events [3, 41, 51]. In order to shed light on these questions, we chose to study the microbial communities of these lakes because, due to their different environmental features, it should be possible to distinguish ecotypes within the same species. We compared OTUs-97% with ASVs and we examined whether ASVs actually corresponded to different environmental variables such as mixing regime, depth layer, water temperature or dissolved oxygen. We also studied the dynamics of these ecotypes across space and time and how their existence affected not only the persistence of species, but also the stability of the whole community itself.

## Materials and methods

### Samples

We analyzed a set of 1,505 samples taken by the K. McMahon laboratory (https://mcmahonlab.wisc.edu/) from eight different temperate bog lakes mainly during the growing seasons of 2005, 2007, 2008, and 2009. Lakes are located in Wisconsin (U.S.A.) (**Fig. 1**) (see [41] for more information). According to their mixing regime, three of the lakes were polymictic, three dimictic, and two meromictic. Samples were taken from the epilimnion as well as from the hypolimnion at different depths.

**Fig. 1.**
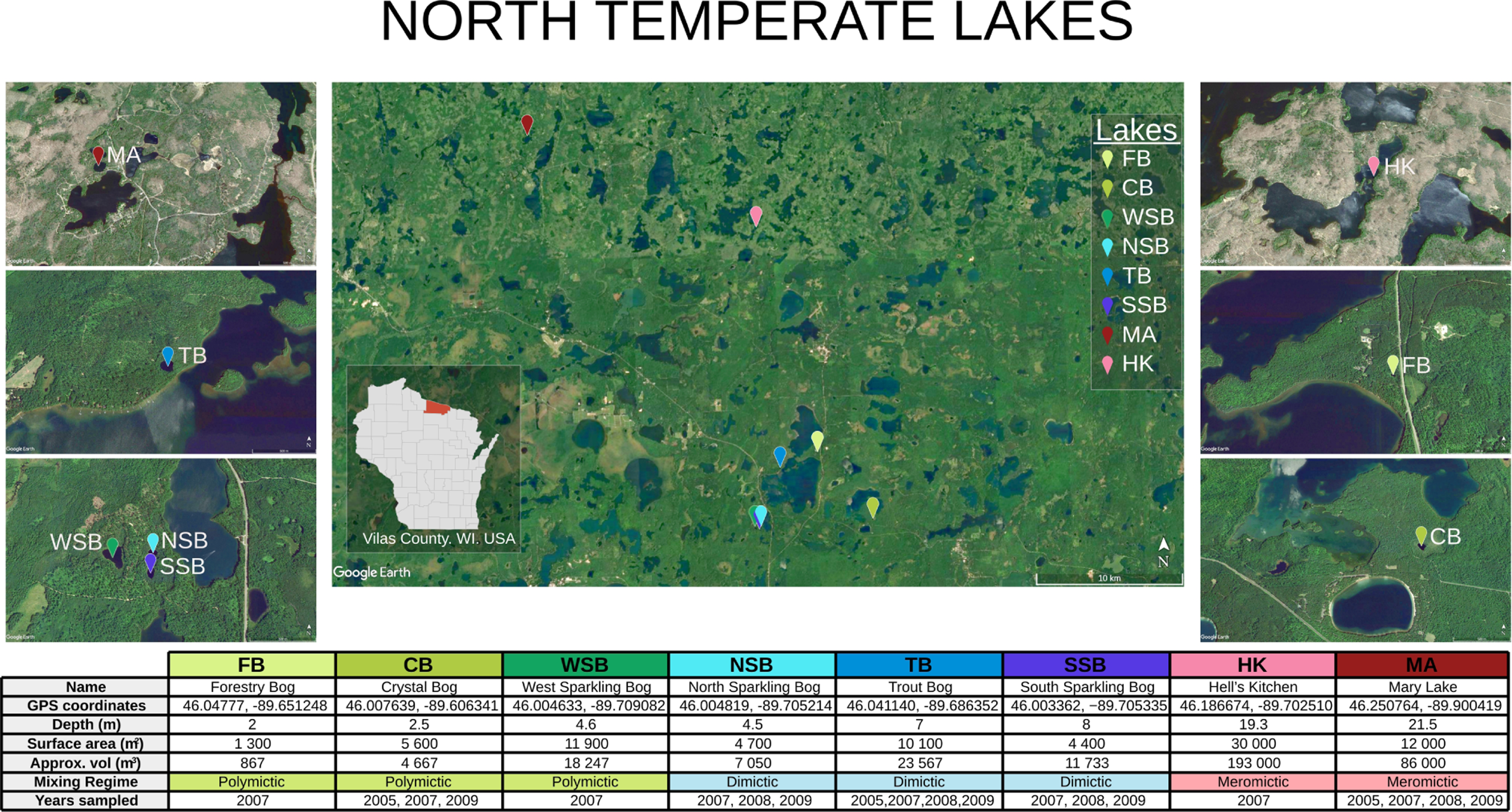
Characteristics of sampling lakes. Table adapted from Linz *et al.* (2017) [41]

Sequence data used in this study were publicly available in the European Bioinformatics Institute with accession number ERP016854 [41] (they can be found also in SRA with the accession number ERS1293140). The metadata linked to these samples were available in the MG-RAST server (mgp127) [52] and in the McMahon Lab GitHub repository “North_Temperate_Lakes-Microbial_Observatory” (https://github.com/McMahonLab/North_Temperate_Lakes-Microbial_Observatory). The 150 bp sequences corresponded to amplicons from the V4 region of the 16S rRNA gene, which was amplified and sequenced using the Illumina HiSeq platform. A complete description of the sample collection and processing protocols can be found in [40, 41].

### High-resolution microbial diversity

We quality-filtered raw sequences using *moira* [53] truncating them to 140 base pairs and discarding every sequence with above 0.005% probability of having more than one error according to its quality scores (*maxerrors*=1, *alpha*=0.005). Then, we processed the sequences with a common pipeline based on the DADA2 software [23] to distinguish real variants, called ‘Amplicon Sequence Variants’ (ASV), from errors. Briefly, the DADA2 pipeline included the following steps: calculation of error rates from sequences, de-replication of amplicon sequencing reads, inference of the composition of the samples, removal of chimeras using the ‘consensus’ method, and assignment of taxonomy using SILVA nr v.128 [54, 55]. We selected bacterial sequences only. Then we removed those samples with less than 10,000 sequences or without associated metadata and rarefied the remaining samples to 10,000 sequences. After this step, we retained 26,128 ASVs from 1,108 samples. These ASV sequences are equivalent to OTUs (Operational Taxonomic Units) that differ in as little as one base out of 140 base pairs, therefore corresponding to more than 99.3% identity. The final result was a table containing the abundance of each ASV in each sample. The sequences were taxonomically classified against the FreshTrain database [45] using mothur v.1.39.1 [56]. More detailed information about the successive steps of the procedure can be found in **Supplementary Table 1S**.

Additionally, we clustered the ASVs at 97% of sequence identity with the Opticlust algorithm [57] in mothur [56]. Thus, we obtained 13,940 OTUs at 97% identity level, and we created a second abundance table for these OTUs-97% (**Supplementary Table 2S**).

### Statistics

We normalized the data with a Hellinger transformation. This transformation is particularly suited to abundance data because it gives low weights to variables (OTUs-97% or ASVs) with low counts and many zeros [58, 59]. The transformation itself comprises dividing each value in the data matrix by its row sum, and taking the square root of the quotient [58, 59]. Then, we did a Detrended Correspondence Analysis (DCA) [60] in terms of similarity distances among samples calculated from the abundance of OTUs-97% or ASVs respectively, separately for the epilimnion and hypolimnion layers. We performed a Procrustes analysis in order to compare both DCAs [61].

Finally, we tested whether the samples clustered significantly by lake calculating the Bray-Curtis distance matrix from the original abundance table and applying a PERMANOVA test (*adonis*). All statistical analyses were done with R version 3.4.4 and the package vegan [62].

### Environmental preferences of ASVs

In order to determine the environmental preferences (lakes and layers) for each ASV, we calculated their mean abundances in each lake-layer combination and standardized them to a mean of zero and a standard deviation of one (z-score). This normalization facilitated visualization of the habitat preference for low-abundance ASVs.

### Influence of intra-species diversity on persistence and stability

The *effective intra-species diversity* was calculated as follows. First, we selected the OTUs-97% with a global abundance higher than 5,000 counts, as the Shannon diversity index has been shown to lose accuracy for abundance values below that threshold [63]. For each OTU-97%, we calculated the global abundances of its constituent ASVs, and rarefied the resulting vector to 5,000 counts. This was done in order to avoid OTUs-97% with higher global abundances being assigned a higher intra-species diversity. Finally, for each OTU-97%, the *effective intra-species diversity* was calculated from the rarefied abundances of its constituent ASVs as the exponential of the Shannon index [64]. This value represents the effective number of ASVs in a given OTU-97%, and its calculation is conceptually analogous to the calculation of the *effective number of species* in a community, but using ASVs as units of diversity and restricting the sampling space to each individual OTUs-97%.

For each OTU-97% with a global abundance above 5,000 and ASV with a global abundance above 1,000, we also calculated its persistence (fraction on samples in which the taxon was present) and variability (coefficient of variation of the abundance of the taxon across the samples), as described in [41]. We then tested for differences in persistence and variability between OTUs-97% and ASVs and between OTUs-97% with low (less than two effective ASVs) and high (two or more effective ASVs) intra-species diversity via Welch’s t tests.

### OTU-97% and ASV distribution modeling

We selected the most stable OTUs-97% (global abundance > 5,000, persistence > 0.75 and variability < 2) and their constituent ASVs (global abundance > 1,000). OTU-97% and ASV distribution in samples was modeled as a function of lake, year, month, depth, temperature, and dissolved oxygen concentration using boosted regression trees as implemented in the *dismo* R package [65]. The *gbm.step* function was used to select an optimal number of boosting trees with the following parameters: *tree.complexity* = 5, *n.trees* = 1,000, *max.trees* = 20,000, *step.size* = 10, *bag.fraction* = 0.75, *learning.rate* = 0.001. The influence of each environmental variable on the distribution of each OTU-97% / ASV was measured as its relative influence on reducing the loss function during model computation. OTUs-97% with only one high abundance ASV and models with a cross-validation correlation smaller than 0.6 were excluded from further analyses.

We tested whether temperature and oxygen influenced the abundance ratios between ASVs belonging to the same clades. For each lake, we selected three significant clades (*Polynucleobacter* PnecC, *Actinobacteria* acI-B2 and *Bacteroidetes* bacI-A, see results for the reasons and **Supplementary Table S3**) and the three most abundant ASVs from each one. For each possible pair of ASVs from the same clade (1-2, 1-3, and 2-3), we then fitted a linear model of their abundance ratio (in log scale) against temperature separately for each lake. The resulting p-values were corrected for multiple testing with the Bonferroni method. For each lake and clade, the two ASVs with a higher correlation between abundance ratio and either temperature or oxygen were selected for further analysis.

## Results

### Diversity and dynamics of the bacterioplankton

First, we analyzed the bacterial diversity in the lakes at the OTU-97% level (**Supplementary Fig.S1**). The most abundant OTUs-97% considering all the lakes belonged to phyla *Proteobacteria, Actinobacteria, Acidobacteria, Bacteroidetes*, and *Verrucomicrobia*. The most prevalent OTU-97% was the betaproteobacterium *Polynucleobacter necessarius* (PnecC), which was very abundant in all the lakes, followed by *Actinobacteria* acI-B2. Lakes with the same mixing regime shared the same abundant taxa through the seasons (**Supplementary Fig. S1**). For instance, *Polynucleobacter* PnecC and *Actinobacteria* acI-B2 were the most abundant OTUs-97% in polymictic and dimictic lakes, but they were accompanied by *Actinobacteria* acI-B3 and *Verrucomicrobia* in polymictic, and *Gammaproteobacteria* and *Acidobacteria* in dimictic lakes. In contrast, these OTUs-97% were less abundant in meromictic lakes.

In order to evaluate the similarity in bacterial composition among the samples, we applied a detrended correspondence analysis (DCA) to both the OTUs-97% and ASVs abundance data, distinguishing between epilimnion and hypolimnion (**Fig. 2**). At both OTUs-97% and ASVs levels, the samples clustered by lake and mixing regime (PERMANOVA significance test p < 0.001, **Supplementary Table S4**). Samples from meromictic lakes were very different from the others, explaining most of the variance among samples. In contrast, samples from the other two mixing regimes showed less differences in bacterial composition. Samples for WSB clustered independently from the other polymictic lakes. This lake is larger in surface area and deeper than the other two polymictic lakes, and this might be the cause of the different community composition.

**Fig. 2.**
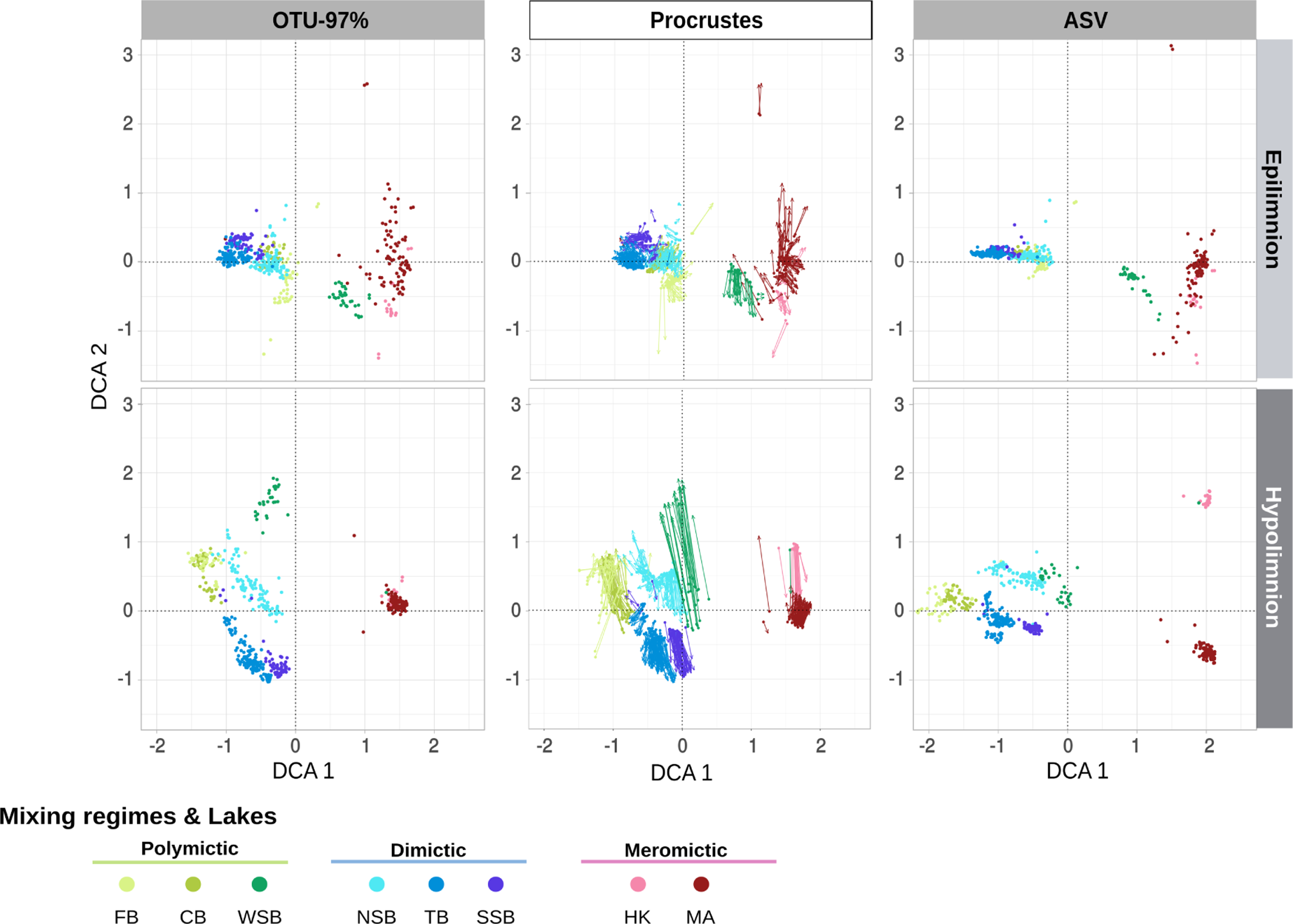
Detrended Correspondence Analysis (DCA) of the bacterial composition in OTUs-97% and ASVs for the epilimnia and hypolimnia of the different lakes. Each point represents a sample. Lakes are indicated by color symbol. Procrustes analysis in the middle column shows the different between the OTU-97% and ASV-level ordinations. Arrow length indicates the difference in the ordination at OTUs-97% and ASVs levels for each sample.

The clustering of samples with OTUs-97% and with ASVs exhibited rather good agreement, as shown by Procrustes analysis (**Fig. 2**), since samples clustered by lake of origin in both cases. Information from ASVs, however, suggested finer distinctions in the sample composition given that, for example, hypolimnion samples from lakes HK and MA clustered together in the OTU-97% analysis, but they were separated when using ASVs along DCA component 2 (**Fig. 2**).

To search for possible seasonal patterns, we also plotted time-decay curves for the four lakes that were sampled for two or more consecutive years (**Supplementary Fig. S2;** Lakes MA, NSB, TB, and SSB). Regardless of whether we considered OTUs-97% or ASVs, Bray-Curtis dissimilarities were maximal for pairs of samples from opposite seasons (i.e. separated 6, 18, 30 months) and minimal for pairs of samples collected at the same time of the year (i.e. separated 12, 24, 36 months). However, even in the latter case, Bray-Curtis dissimilarities were relatively high (0.521 for ASVs and 0.462 for OTUs-97%), showing that, albeit seasonal trends could be discerned, there was a high inter-year variability in community composition.

### ASVs from the same OTU-97% show different habitat preferences

In order to explore the ecological significance of intra-species diversity, we analyzed the distribution and habitat preferences of the ASVs corresponding to the most abundant OTU-97% (**Fig. 3**). The upper row of radar plots in **Fig. 3** shows the average abundance of ASVs from each of the most abundant OTUs-97% in the different lakes (color coded inner ring) and layers (grey scale coded outer ring).*Polynucleobacter* PnecC was the most abundant OTU-97% in the dataset, comprising three main ASVs (ASV-1, ASV-2 and ASV-48). ASV-1 was more abundant than ASV-2 in the hypolimnion of NSB, but less abundant in the epilimnion of FB. Both ASVs, in turn, were always more abundant than ASV-48.

**Fig. 3.**
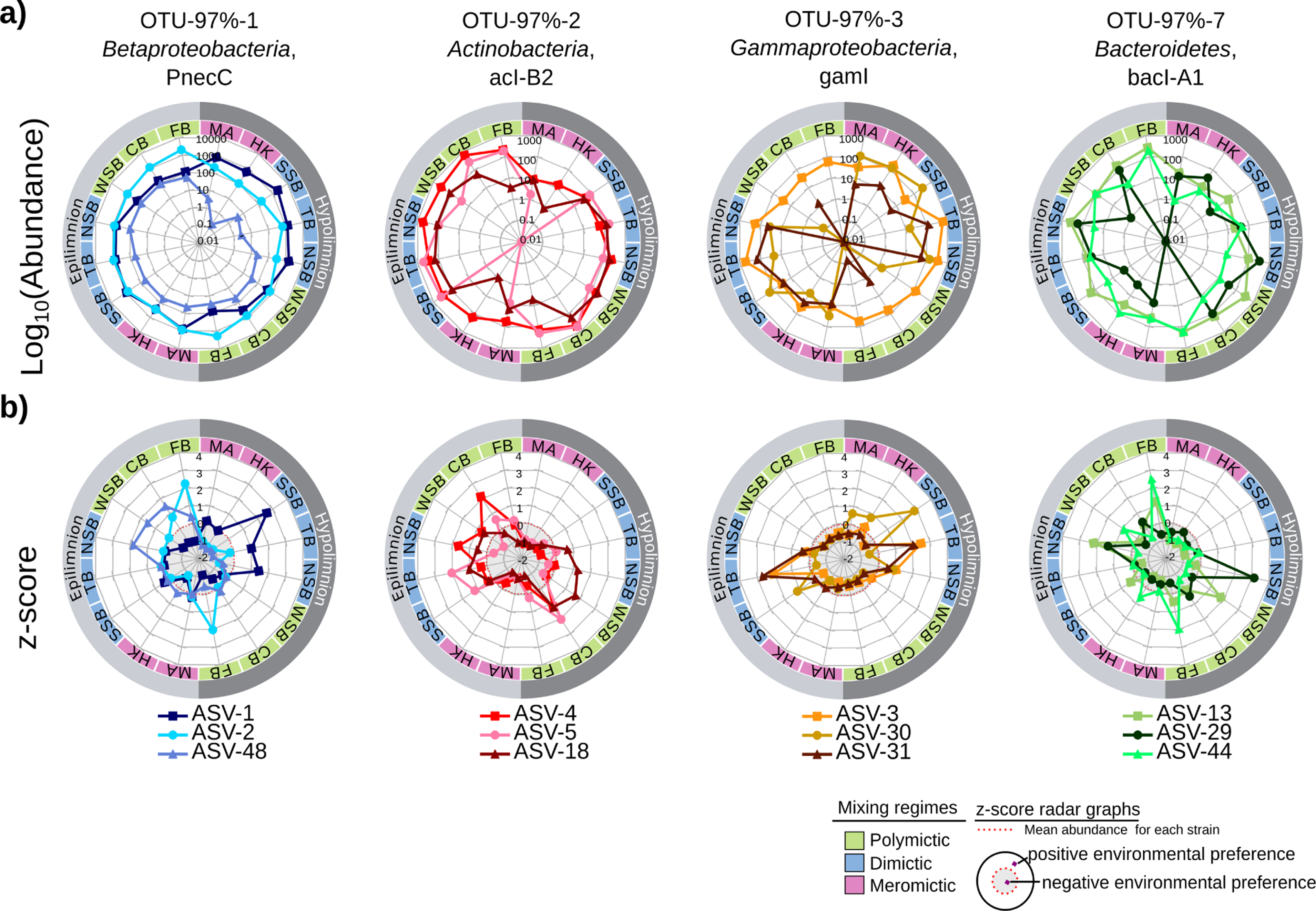
Environmental preferences. Distribution and habitat preference of the four selected OTUs-97%. The radar plots show the abundance (logarithmic scale, a) and z-score (b) of the three most abundant ASVs for each of four OTUs-97%. The abundances of the ASVs inform about their real distribution. The colors of the inner ring represent the mixing regime of the lake: green for polymictic, blue for dimictic and red for meromictic. The shades of grey in the outer ring indicate the lake layer (epilimnion and hypolimnion). Zero values have been plotted as log10 (0.01). z-scores indicate the environmental preference of each ASV. Since the z-score normalization was made independently for each ASV, z-scores values cannot be compared quantitatively among ASVs. Red dotted line in z-score plots indicate the mean of their abundances, separating positive (white background) and negative (grey background) environmental preferences as schematically represented at the bottom right of the figure. ASV’s points above the dashed red line exhibit a positive preference for that environment and vice-versa.

In order to determine the habitat preference of ASVs irrespectively of their abundance, z-scores were plotted in the lower row of radar plots (**Fig. 3**). Positive z-scores represent lake/layer samples in which the ASV had higher-than-average abundance, and vice-versa. The three most abundant ASVs from *Polynucleobacter* PnecC showed different preferences: ASV-1 showed a higher abundance in the hypolimnion of dimitic lakes, ASV-48 in the epilimnion of polymitic lakes, and ASV-2, despite being present in most lakes, was most abundant in FB. Differences in the distribution of ASV-1 and ASV-2 in different lakes can be seen in **Supplementary Fig. S3**. *Polynucleobacter* was present in all lakes, while ASV-2 was prevalent in polymictic lakes. ASV-1 and ASV-2 alternated their abundance in the remaining lakes.

Similar situations were found for the other OTUs-97%. In the case of *Actinobacteria* acI-B2, ASV-4 preferred CB, while ASV-18 tended to appear in the hypolimnion of dimictic and polymictic lakes. ASV-5, on the other hand, showed avoidance of the meromictic lakes, especially of HK, but it showed some preference for several epilimnetic samples and the hypolimnion of CB. The *Gammaproteobacteria* gamI (which belongs to the genus *Methylobacter*) illustrates a case in which two strains of the same species had similar spatial preferences: ASV-3 and ASV-31 both preferred TB. However, ASV-31 was absent from several lakes where ASV-3 was present. In *Bacteroidetes* bacI, finally, ASV-29 preferred CB and NSB, while ASV-44 was more abundant in FB (**Fig. 3**).

We found similar situations in less abundant OTUs-97%. For instance, *Acidobacteria Geothrix* (the fifth most abundant OTU-97%) was present in lakes TB and SSB (**Supplementary Fig. S1**), but looking at the ASV level, ASV-6 was more abundant in TB while ASV-15 in SSB (**Supplementary Fig. S3, Supplementary Fig. S4**). *Actinobacteria* acI-B3 was present in the three polymictic lakes (**Supplementary Fig. S3)**. ASV-8 was dominant in FB but both ASV-8 and ASV-10 co-dominated in the other two lakes (**Supplementary Fig. S3, Supplementary Fig. S4**). In general, abundant OTUs-97% were present in most lakes and layers, but their constituent ASVs showed preferences for only some of them. Our hypothesis is that such distributions of ASVs reflect slightly different adaptations to the environmental conditions in each layer and lake.

### ASVs from the same OTU-97% show different temporal dynamics

ASVs from the same OTU-97% not only showed different spatial distributions, but also different temporal dynamics, as exemplified in NSB (**Fig. 4**). For instance, *Polynucleobacter* PnecC had a fairly constant relative abundance during the 2007-2008 period (**Fig. 4a**). However, its two most abundant ASVs (ASV-1 and ASV-2) changed their abundances over time: in general, ASV-1 was more abundant in the hypolimnion and ASV-2 in the epilimnion, but ASV-1 came to dominate also in the epilimnion during Spring 2008, when the surface water was still cold (**Fig. 4a**). Similar trends were observed in the dimictic lakes TB and SSB (**Supplementary Fig. S3)**. This suggests that each ASV may be adapted to different water temperatures.

**Fig. 4.**
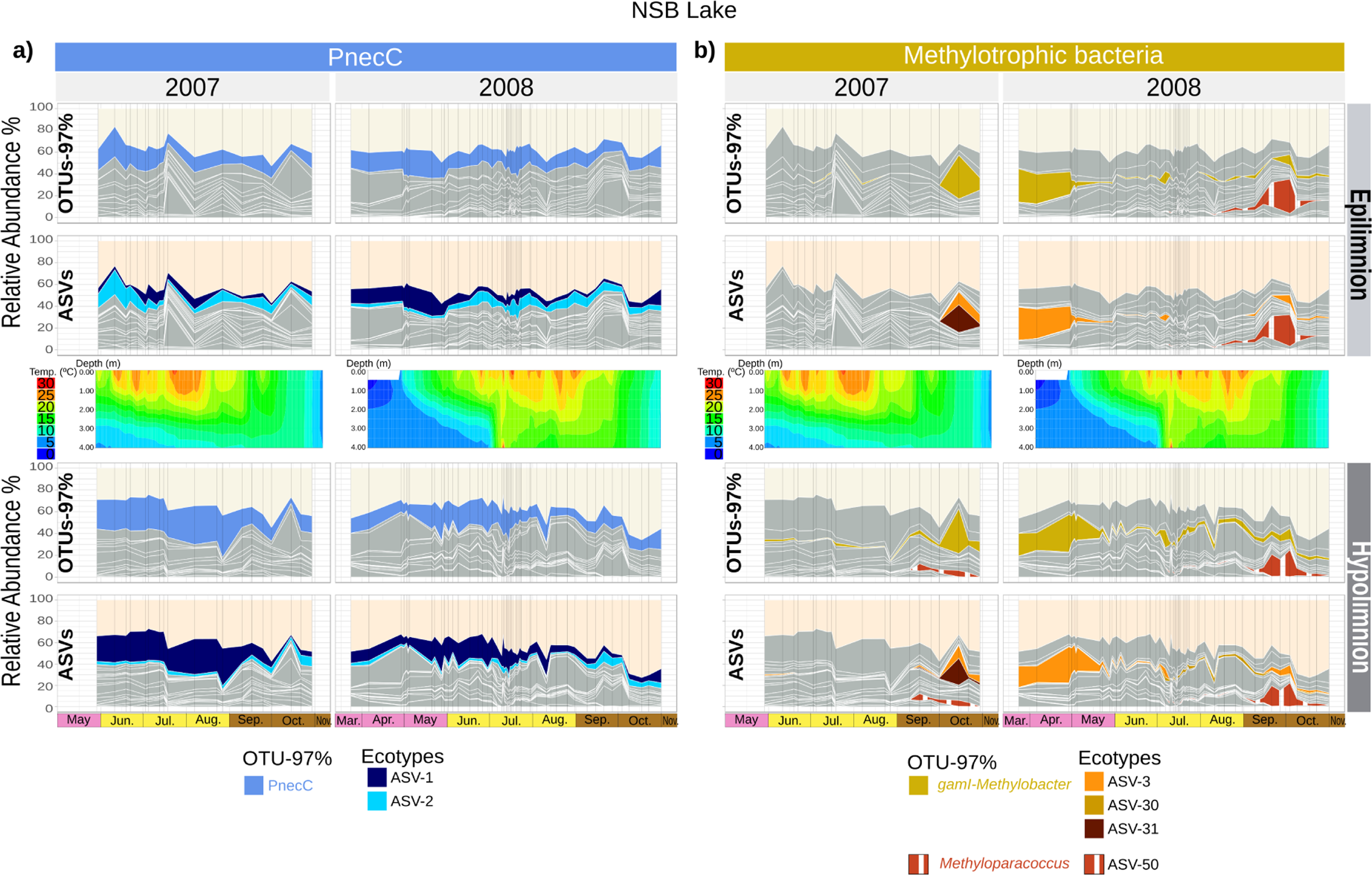
Microdiversity shows temporal patterns in NSB. **a)** Temporal abundance pattern of PnecC OTU-97% and its corresponding ecotypes with time. **b)** Temporal abundance pattern of gamI OTU-97% and their corresponding ecotypes with time. In both cases, the middle row shows the temperature profiles along time. Homogeneous vertical temperature profiles correspond to mixing events. In 2008, there was an artificial mixing event from July 2 to July 10, when homogeneous temperature was observed. Black vertical lines in the different panels indicate sampling points.

ASV resolution also provided new insights into the behavior of species experiencing episodic blooms. *Gammaproteobacterium* gamI, which could be assigned to the methanotrophic genus *Methylobacter*, bloomed during the fall mixing of 2007, remained abundant until the spring mixing, and dropped to a very low abundance during summer stratification (**Fig. 4b**).

This abundance pattern is consistent with the combined need of methane (produced in the sediments by methanogens) and oxygen (exchanged with the atmosphere at the surface). During summer stratification, methane and oxygen would only be found together in the thermocline separating the epilimnion and the hypolimnion, hence the low abundance of gamI. Due to the lack of mixing, methane could neither be consumed in the hypolimnion (as there is no oxygen available) nor released to the upper layer (save for diffusion, which is a comparatively slow process [66]). Thus, methane would slowly build up in the hypolimnion during summer.

During the fall mixing, however, hypolimnetic methane would be mixed with epilimnetic oxygen throughout the whole water column, leading to the observed bloom of gamI in 2007 (**Fig. 4b**). This bloom was followed by a sharp decrease in oxygen concentrations (**Supplementary Fig. S5**, [3]), which we attribute to the rapid oxidation of the methane burst by gamI. This methanotroph was actually represented by two different ASVs in NSB: ASV-3 and ASV-31. ASV-31 was only abundant at the beginning of the autumn mixing event, concurrent with the decrease in oxygen concentrations.ASV-3, on the other hand, dominated afterwards, when oxygen concentrations returned to higher levels (**Fig. 4b**, **Supplementary Fig. S5**, [3]).

We hypothesize that the two ASVs represent two ecotypes of *Methylobacter*, with ASV-31 being adapted to compete for oxygen when there is an excess of methane, and ASV-3 being suited to low methane / high oxygen scenarios. This would be consistent with the known existence of *Methylobacter* groups with different oxygen preferences, some of which are able to live at undetectable concentrations of oxygen [67]. Interestingly, in the fall mixing of 2008 a different methanotrophic OTU-97% (*Methyloparacoccus*, ASV-50) bloomed at about the same time (**Fig. 4b**). These data point to a double layer of functional redundancy: there were two OTUs-97% carrying out methanotrophy and blooming after fall mixing in different years, and at least two ASVs with different temporal dynamics within *Gammaproteobacteria* gamI.

### Intra-species diversity imparts stability at the species level

In general, all abundant OTUs-97% were represented by at least two ASVs with different distributions in space and seasonal dynamics. This points to the existence of different ecotypes with slightly different environmental preferences. We explored whether rich intra-species diversity was linked to increased stability at the species level. To that end, we calculated the effective intra-species diversity of the OTUs-97% from the abundances of their constituent ASVs (see materials and methods). We found out that OTUs-97% with two or more effective ASVs were more persistent (i.e. appeared in a larger proportion of samples) and less variable (i.e. showed a more constant abundance across samples) than OTUs-97% with less than two effective ASVs (Welch’s t test, p < 0.001 in both cases, **Fig. 5**).

**Fig. 5.**
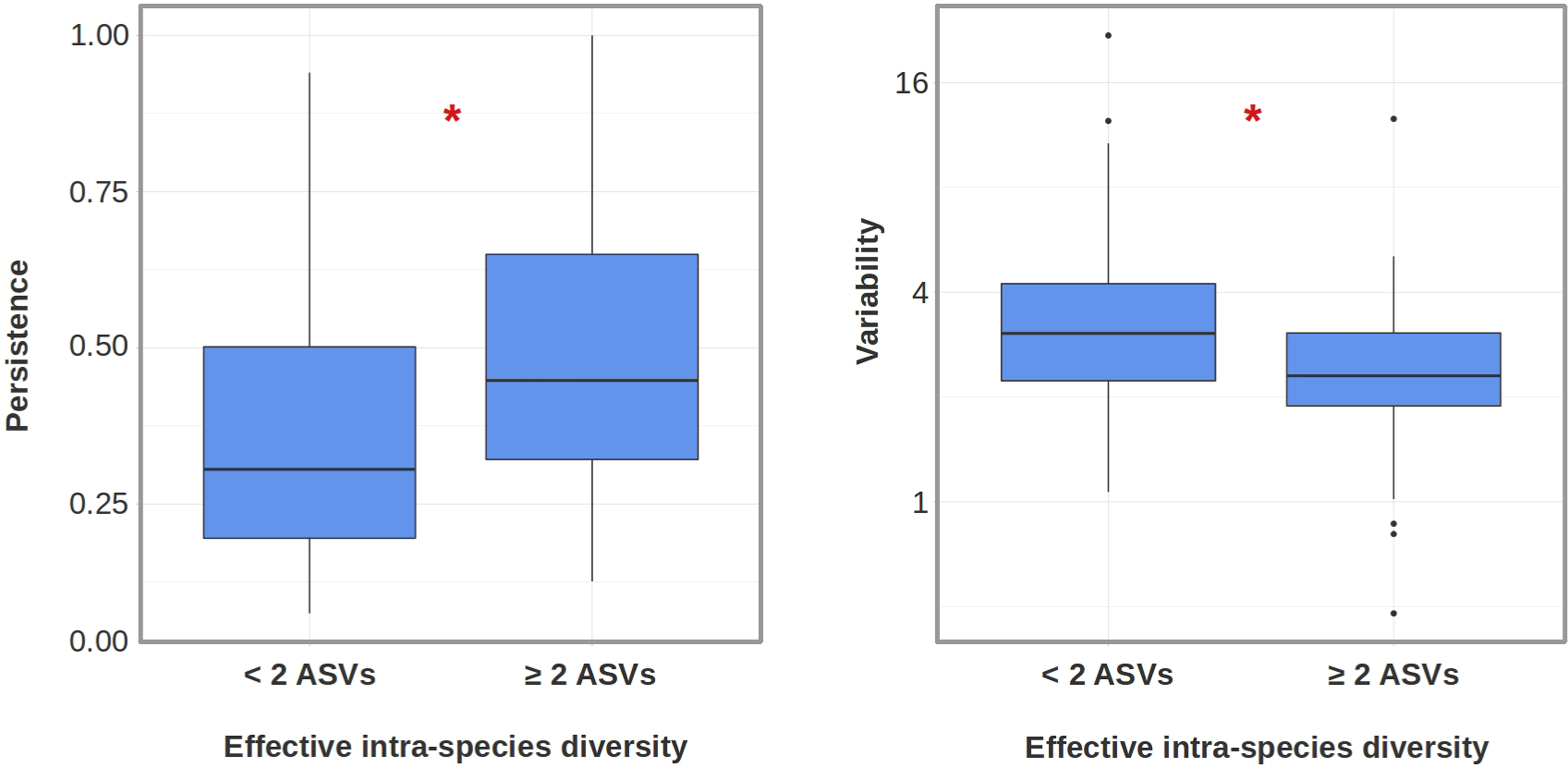
Effect of intra-species diversity on species persistence (left) and variability (right). Boxplots show the persistence and variability of OTUs-97% with low (< 2 effective ASVs) and high (2 or more effective ASVs) effective intra-species diversity. Significant (Welch’s t test *p* < 0.001) differences are denoted with an asterisk.

### Environmental factors drive intra-species diversification

We were interested in finding whether intra-species diversification occurred in response to a particular environmental gradient. To test this, we modeled the distribution of the most stable OTUs-97% (persistence > 0.75, variability < 2) and their constituent ASVs across samples as a function of lake, year, month, depth, temperature, and dissolved oxygen concentration, using boosted regression trees [65]. The median cross-validation correlation of our models was 0.79, indicating that OTUs-97% and ASV abundances in the North Temperate Lakes could be reasonably modeled from the available parameters. We then compared the contribution of the different environmental parameters to the OTU-97% models and the ASV models (**Supplementary Fig. S6**). Temperature and oxygen concentration had a significantly higher influence on the distribution of individual ASVs than on the distribution of their parent OTUs-97% (Welch’s t test p = 0.041 and p = 0.014, respectively), suggesting that intra-species diversification in response to temperature and oxygen gradients is a general strategy in these freshwater lakes.

### Dynamics of a freshwater model community

In order to identify potential associations between the taxa present in our samples, we further focused on representatives from a freshwater model community identified by Garcia *et al*. [68]. This community was originally enriched from the dimictic Lake Grosse Fuchskuhle, and consisted of four organisms commonly found in freshwater environments (*Polynucleobacter* PnecC, two different *Actinobacteria* and a *Bacteroidetes*), which were able to grow together in a mixed culture presumably by establishing mutually beneficial metabolic interactions ([68]; **Fig. 6a**). Even though each strain presented different auxotrophies that precluded them from growing alone, the consortium was stable over time [68], thus revealing a powerful model of mutualism in natural microbial communities.However, as noted by the authors, mixed cultures provide limited information on how the system may evolve under changing environmental conditions [68]. We therefore sought to identify members of that model community in our dataset, and to characterize their dynamics over time.

**Fig. 6.**
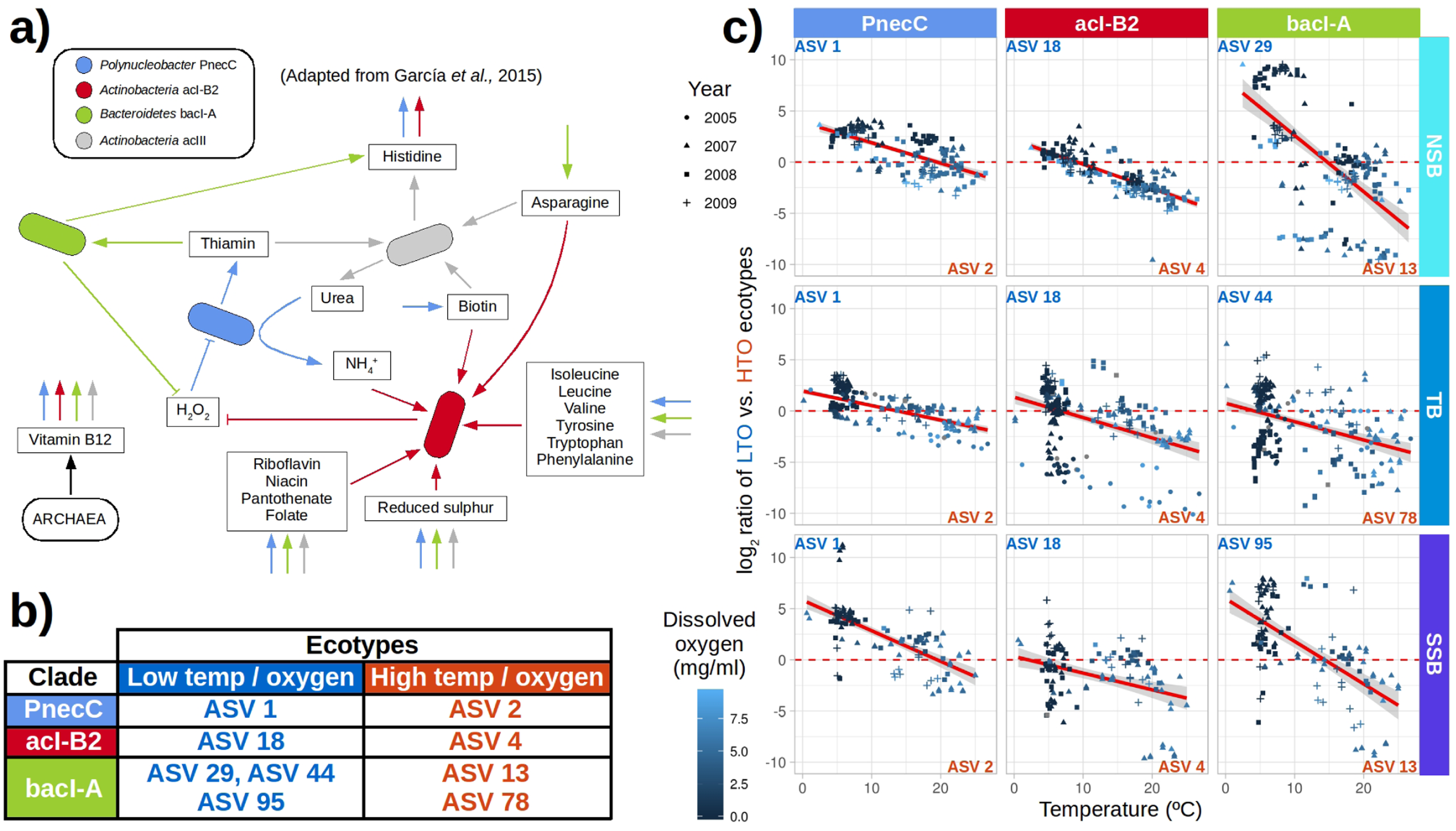
Microdiversity provides community stability across environmental gradients in freshwater ecosystems. **a)** Freshwater model community proposed by García *et al.* (2015) [68]. **b)** Amplicon sequence variants (ASV) from the different member clades of the model community, classified according to their environmental preferences. **c)** Relative abundance of low temperature/oxygen (LTO) versus high temperature/oxygen (HTO) ecotypes of the model community in dimictic lake samples, plotted as a function of water temperature. Symbol color indicates dissolved oxygen concentration in shades of blue. Symbol shape indicates sampling year. Samples above the dashed red line are dominated by LTO ecotypes, and vice-versa. Significant linear regressions (FDR < 0.05) and 95% confidence intervals are shown with a red line and a shaded grey area, respectively.

Of the four members, *Polynucleobacter* PnecC, actinobacterium acI-B2 and a bacteroidetes from the *Cytophagaceae*/bacI-A clades were found in high abundances in our data, in particular during the summer season. Conversely, actinobacterium acIII was either absent or present in very low abundance in almost all samples. This might be due to acIII being more successful when growing in culture than in the environment, or else to differences between Lake Grosse Fuchskuhle and the North Temperate Lakes analyzed in this study. Free-living freshwater bacteria have been recently proposed to establish promiscuous rather than specific interactions, fulfilling their metabolic needs by associating with whatever suitable partner they happen to encounter [69]. The ecosystem role of acIII might thus be fulfilled by a different microorganism in the North Temperate Lakes.

The PnecC, acI-B2, and bacI-A clades were represented by more than one ASV in our dataset, and the abundance of each ASV was largely controlled by temperature and oxygen. The most abundant ASVs for each clade and each of the dimictic lakes are shown in **Fig. 6b**, together with their temperature and oxygen preferences. Both parameters significantly explained the abundance ratios of the different strains from the same clade in the three dimictic lakes analyzed in this study (**Fig. 6c**, FDR < 0.05), as well as in most of the other lakes (**Supplementary Fig. S7**). Oxygen concentrations appeared as a secondary factor, as the example of bacI in Lake WSB illustrates (**Supplementary Fig. S8**). For the most part, ecotypes appeared to respond similarly to temperature and oxygen concentrations, which were in fact significantly correlated in our data. The only exceptions were the two main ecotypes of clade bacI in NSB (dependent on oxygen, but not on temperature) and of clade acI-B2 in SSB (dependent on temperature, but not on oxygen). The existence of niche partitioning among ecotypes of the same species has been reported previously [70–73], and our results suggest that this phenomenon might be generalized in freshwater ecosystems.

## Discussion

The comparative study of eight temperate lakes of different characteristics has provided insights on the microbial ecology of these systems. We first focused in evaluating whether OTUs-97% and ASVs offered different perspectives on the compositional dynamics of freshwater lakes. Ordination methods showed a good agreement between the OTU-97% and ASV resolutions in explaining the general compositional trends of our samples (**Fig. 2**), suggesting that community structure was at least partially determined at the species, rather than the ecotype, level. This agrees well with previous reports of ASV and OTU-97% methods revealing similar broad scale ecological results [74]. Samples from the same lakes and different years clustered together (**Fig. 2**), showing that each lake had a characteristic compositional signature that remained relatively stable over time.

In spite of this, there was a high temporal species turnover in the four lakes that were sampled in consecutive years (**Supplementary Fig. S2**): although seasonal patterns could be observed, the communities did not return to their exact original composition after a year had passed. Freshwater microbial communities are composed by metabolically streamlined microorganisms, which thrive by promiscuously cooperating with a wide array of possible partners [69]. This versatility should result in a compositionally fluid microbial community, which may remain robust without necessarily showing strong seasonal trends. The lake NSB, for example, hosted blooms associated with the fall mixing in both 2007 and 2008, but the blooms originated from two different species (a *Methylobacter*-like OTU-97% in 2007, and a *Methyloparacoccus*-like OTU-97% in 2008; **Fig. 4b**). Both species are known methanotrophs, which makes it likely that their respective blooms responded to the same underlying environmental trigger (the combined availability of hypolimnetic methane and epilimnetic oxygen during the mixing event) and had a similar effect on the ecosystem (a transient depletion of oxygen associated with methane consumption, **Supplementary Fig. S5**). In this sense, even if community composition changed from year to year, the associated functional turnover may have been much smaller.

Despite this interannual variability in community composition, some core taxa were found in all communities and layers, as previously described in [41]. When we focused in the most abundant OTUs-97%, we found that they were all composed by different ASVs, with different preferences for particular lakes and layers (**Fig. 3**). This hinted at an important role of intra-species diversity in the dynamics of the microbial communities of lakes. While it is true that OTUs-97% and ASVs provided a similar overview of community structure (**Fig. 2**), some ecosystem processes might be determined by the presence and interplay of particular ecotypes, which can be only distinguished by delineating ASVs [19–22].

Several studies have described the existence of ecotypes within microbial species in several environments such as host-associated microbiomes [20–22, 28, 29, 75], granular biofilms [76], marshes [22], the ocean [12, 34, 77, 78], oil spills [30, 31] or freshwater lakes [79–82]. Moreover, it has been claimed that a rich biodiversity guarantees the long-lasting maintenance of bacterial populations [33, 39, 83, 84]. However, the role of intra-species diversity as a stabilizing force in microbial communities remains insufficiently explored. In this study, intra-species diversity appeared to be crucial at preserving the species-level community composition in the face of environmental change. OTUs-97% with more ASVs were more stable than species with one or few ASVs, showing higher persistence and reduced variability (**Fig. 5**). Temperature and oxygen exerted a greater influence on the distribution of ASVs than on the distribution of their parent OTUs-97% (**Supplementary Fig. S6**). This suggests that both environmental factors drive intra-species diversification in ubiquitous freshwater taxa. Indeed, when studying the dynamics of a model community in our samples, we found that three of the most ubiquitous freshwater clades (*Polynucleobacter* PnecC, *Actinobacteria* acI-B2 and *Bacteroidetes* bacI-A) were each represented by two main ecotypes with distinct temperature and oxygen preferences (**Fig.6c**, **Supplementary Fig. S7, Supplementary Fig. S8**). Seasonal changes drove ecotype turnover at both the hypolimnion and the epilimnion, but the community remained consistently represented at the species level.

Our results point at microdiversity as an important factor determining community stability: fluctuations over time (seasonal patterns) and space (mixing layer) may affect the ratio between different ecotypes, but the species in the community remain detectable throughout the year. This is exemplified in **Fig. 4a**, in which the relative abundance of *Polynucleobacter* PnecC was fairly constant during years 2007 and 2008 in both mixing layers, save for blooms of other species during the autumn mixing events. This stability was possible due to the existence of two well-differentiated ecotypes of PnecC (ASV-1 and ASV-2), which replaced each other as environmental conditions changed. In particular, ASV-1, which for the most part was relegated to the hypolimnion (**Fig. 3**; **Fig. 4a**), came to dominate in the epilimnion during early spring, where temperatures where lower and a thermal inversion was still in place. In general, it was difficult to separate the effects of temperature and oxygen, as most of our samples were taken during summer, where water is hotter and more oxygenated in the surface than in deeper locations. However, in early spring, temperature and oxygen concentrations followed opposite trends (**Supplementary Fig. S5**). Thus, the dynamics of both ecotypes of *Polynucleobacter* PnecC during this period suggest that their abundance is controlled by temperature, rather than by oxygen concentrations. This would be consistent with an increasing body of literature highlighting the critical role of temperature in shaping microbial ecosystems [85–87], and in particular, controlling the distribution of *Polynucleobacter* ecotypes [79, 88]. Other authors, however, have pointed at oxygen as a main factor driving *Polynucleobacter* diversification [72], both options are in fact not mutually exclusive, and in any case would result in an increased persistence of *Polynucleobacter* under temporal and spatial variability.

For simplicity, we have assumed that ecotypes from the same species are functionally equivalent, and that their relative abundances in our samples were only controlled by environmental factors. The observed trends in community composition were consistent with such a basic model (**Fig. 6c**), and while this study presents a taxonomic, rather than functional, characterization of our samples, we find it reasonable to assume that phylogenetically close organisms will have similar functional characteristics [87, 89]. Our observations thus fit well with a scenario of widespread functional redundancy [39, 90], with intra-species diversity guaranteeing the persistence of species – and therefore functional stability - across environmental gradients. The idea of functional redundancy has been recently challenged by using OTUs-97% as the units of diversity [91]. As shown in this study, that resolution fails to capture the complex dynamics occurring at the sub-species level, which seem to be widespread - at least in freshwater lakes - and some of which might contribute to functional redundancy [76].

Freshwater lakes are complex ecosystems that host highly diverse microbial communities and have a major role in the global biogeochemical cycles [92–95]. They are also good candidates for studying bacterial community dynamics [3, 41, 96]: they experience seasonal variations in environmental conditions, are subjected to periodic perturbations (mixing), and contain strong environmental gradients (stratification). Lakes have been traditionally believed to show highly seasonal patterns with regards to community composition [79, 97–99], although some multi-year studies have recently partially challenged this view, in particular during the summer season [41]. Lacustrine microbial communities also show variability at short timescales, being sensitive to high-frequency/high-locality perturbations, such as hourly changes in wind speed [100]. Under such unpredictable conditions, adaptability becomes of capital importance for lake-dwelling microorganisms. As recently reported, cooperative promiscuity might be a way to maximize survival within an ever-changing biotic landscape [69]. We now show that intra-species diversification is another factor leading to species persistence, in this case across abiotic gradients. In our data, intra-species diversity was not exclusive of a few taxa, but was a general feature of most from all lakes. We thus believe that widespread intra-specific diversification might be a key process contributing to microbial community stability and ecosystem robustness. While in this study we have mainly focused on dichotomic relationships between ecotypes of the same species, much more complex dynamics might be unraveled in the future. Ongoing long-term time series experiments characterizing not only the taxonomic but also the functional composition of the North Temperate Lakes-Microbial Observatory will surely help to elucidate the links between abiotic and biotic variability, inter and intra-specific diversity, and ecosystem functioning, shedding further light on the way in which microbial communities deal with environmental changes.

## Conflict of interest

The authors declare no conflict of interest.

## Acknowledgments

We thank Guillermo Martín Serrano for his contribution during early stages of data analysis. This work was funded by grant CTM2016-80095-C2-1-R from the Spanish Ministerio de Economía, Industria y Competitividad. N G-G was funded by a student scholarship from the Severo Ochoa Program at CNB (SEV-2013-0347-17-2).

