## Supplementary Information for "Intra-species diversity ensures the maintenance of functional microbial communities under changing environmental conditions"

### Supplementary Material

**Table S1. General information of the pipeline.** Table describing the initial number of sequences and the remaining number of sequences after the analysis.

| | Samples | Sequences | Length (mean) |
|---|---|---|---|
| Start | 1 505 | 67 484 485 | 144.966 |
| After moira (trim, quality filter, no ambiguidities) | 1 505 | 50 755 019 | 140 |
| DADA2-after chimeras | 1 505 | 49089 840 | 140 |

Then Select Bacteria sequences, remove ‘Mitochondria’ and ‘Chloroplast’. Rarefy to n 10 000:

Samples: 1 108
Total Seqs: 11 080 000
Total ASV: 26 128
% Singletons:  5 942 / 26 128 * 100 = 22%
% < 10 counts in all the samples: 17 924 / 26 128 * 100 = 68 %

**Table S2. Summary of ASVs.** Table describing the taxonomy of the ASVs, the OTU-97% they belong to, their sequence and their total abundance in each combination lake-layer.

**Table S3. Taxonomic classification for the members of the studied community.** Table describing the main ASVs for three significant clades in the lakes microbial diversity.

**Table S4 PERMANOVA test.** Table describing if the mixing regimes and the lake are significant factors that influence microbial composition.

**PERMANOVA test**

| | | | Df | Sum of Sqs | Mean Sqs | F. Model | R2 | Pr >(F) |
|---|---|---|---|---|---|---|---|---|
| EPILIMNION | ASVs | **Lake** | 7 | 59.093 | 8.4419 | 54.738 | 0.41914 | 0.001 *** |
| | | **Residuals** | 531 | 81.892 | 0.1542 | | 0.58086 | |
| | | **Total** | 538 | 140.986 | | | 1 | |
| | | **Mixing regime** | 2 | 33.694 | 16.8471 | 84.164 | 0.23899 | 0.001 *** |
| | | **Residuals** | 536 | 107.291 | 0.2002 | | 0.76101 | |
| | | **Total** | 538 | 140.986 | | | 1 | |
| | OTUs-97% | **Lake** | 7 | 44.762 | 6.3945 | 52.853 | 0.41064 | 0.001 *** |
| | | **Residuals** | 531 | 64.244 | 0.121 | | 0.58936 | |
| | | **Total** | 538 | 109.006 | | | 1 | |
| | | **Mixing regime** | 2 | 27.484 | 13.7421 | 90.354 | 0.25214 | 0.001 *** |
| | | **Residuals** | 536 | 81.522 | 0.1521 | | 0.74786 | |
| | | **Total** | 538 | 109.006 | | | 1 | |
| HYPOLIMNION | ASVs | **Lake** | 7 | 89.729 | 12.8185 | 93.318 | 0.54836 | 0.001 *** |
| | | **Residuals** | 538 | 73.902 | 0.1374 | | 0.45164 | |
| | | **Total** | 545 | 163.631 | | | 1 | |
| | | **Mixing regime** | 2 | 53.437 | 26.7183 | 131.66 | 0.32657 | 0.001 *** |
| | | **Residuals** | 543 | 110.194 | 0.2029 | | 0.67343 | |
| | | **Total** | 545 | 163.631 | | | 1 | |
| | OTUs-97% | **Lake** | 7 | 68.396 | 9.7708 | 87.17 | 0.53144 | 0.001 *** |
| | | **Residuals** | 538 | 60.304 | 0.1121 | | 0.46856 | |
| | | **Total** | 545 | 128.7 | | | 1 | |
| | | **Mixing regime** | 2 | 45.551 | 22.7756 | 148.74 | 0.35393 | 0.001 *** |
| | | **Residuals** | 543 | 83.149 | 0.1531 | | 0.64607 | |
| | | **Total** | 545 | 128.7 | | | 1 | |

Permutation: free
Number of permutations: 999
Signif. codes:  0 ‘***’ 0.001 ‘**’ 0.01 ‘*’ 0.05 ‘.’ 0.1 ‘ ’ 1

| | |
|---|---|
| Df | degrees of freedom |
| Sum of Sqs | sequential sums of squares |
| Mean Sqs | mean squares |
| F. Model | F statistics |
| Pr >(F) | partial R-squared and P values |

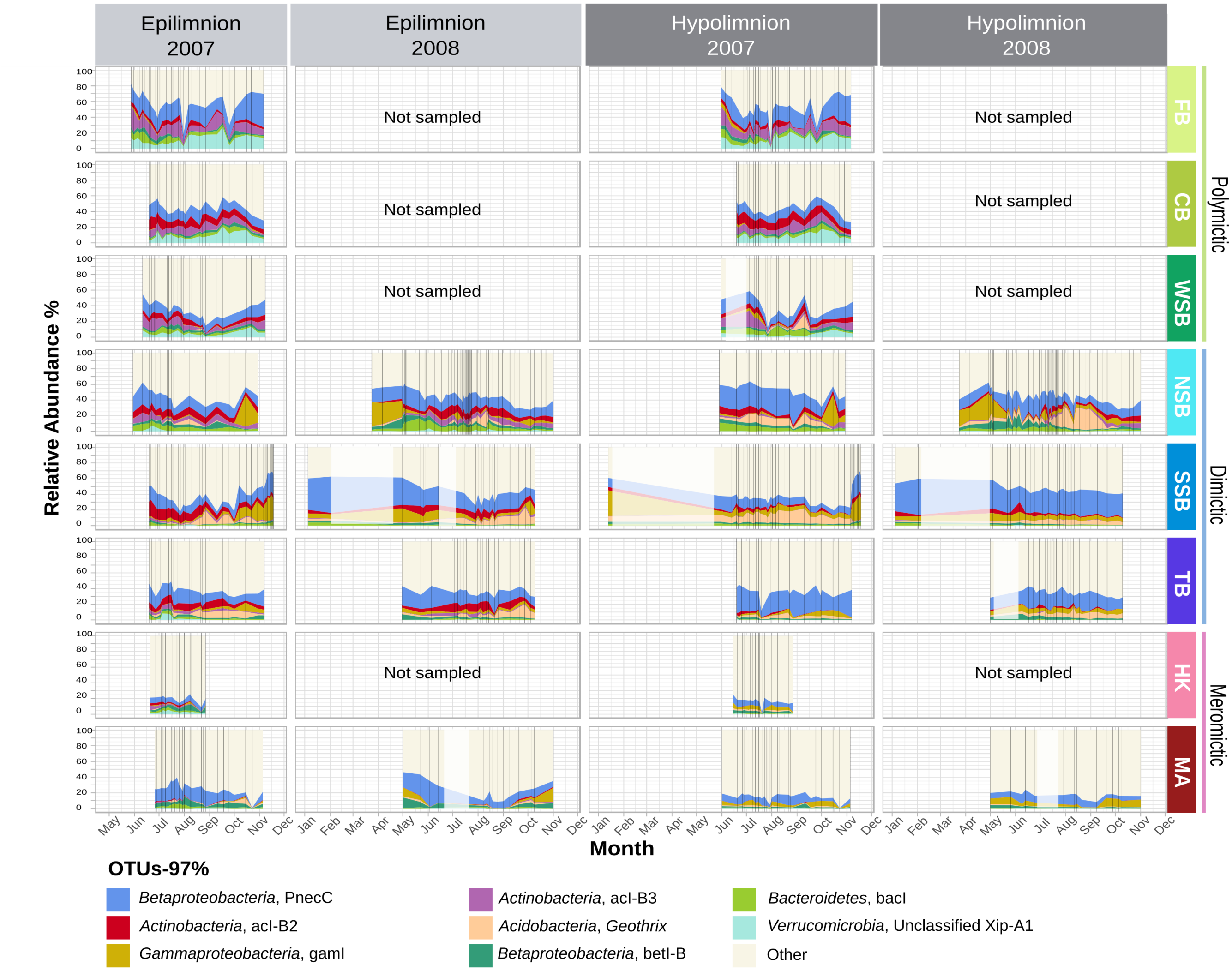


**Fig. S1. Abundance of the eight most abundant OTUs-97% in the dataset.** Vertical lines indicate sampling points. We used linear interpolation between samplings. Periods interpolated longer than 30 days are masked by a white rectangle. The category 'Other' included the sum of the abundances of the OTUs-97% that were not plotted individually.

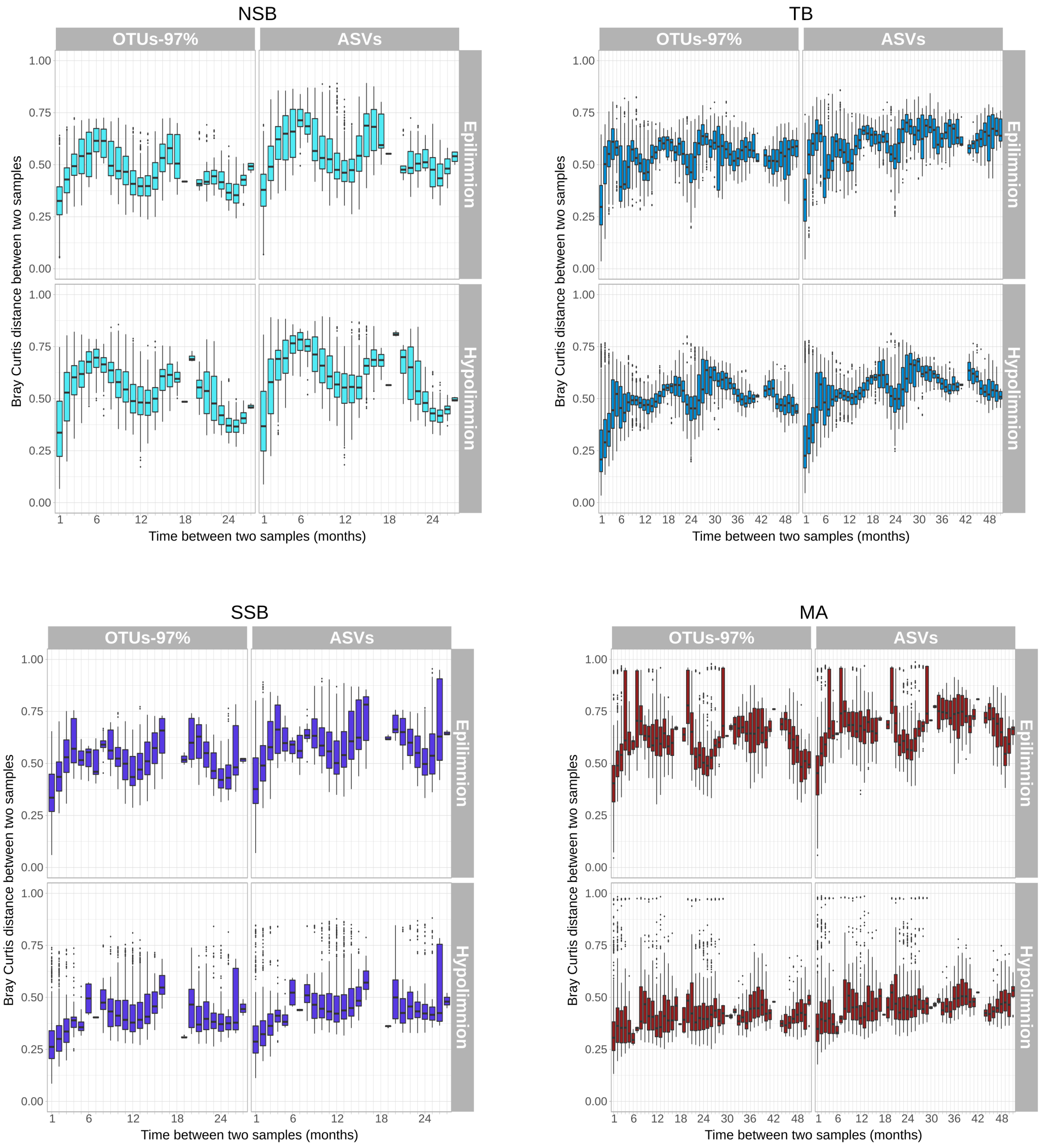


**Fig. S2. Similarity decay for lakes NSB, TB, SSB, MA.** Similarity in community composition for the lakes sampled for two or more consecutive years. Pairwise comparisons of the overall composition for communities using Bray-Curtis dissimilarity are shown.

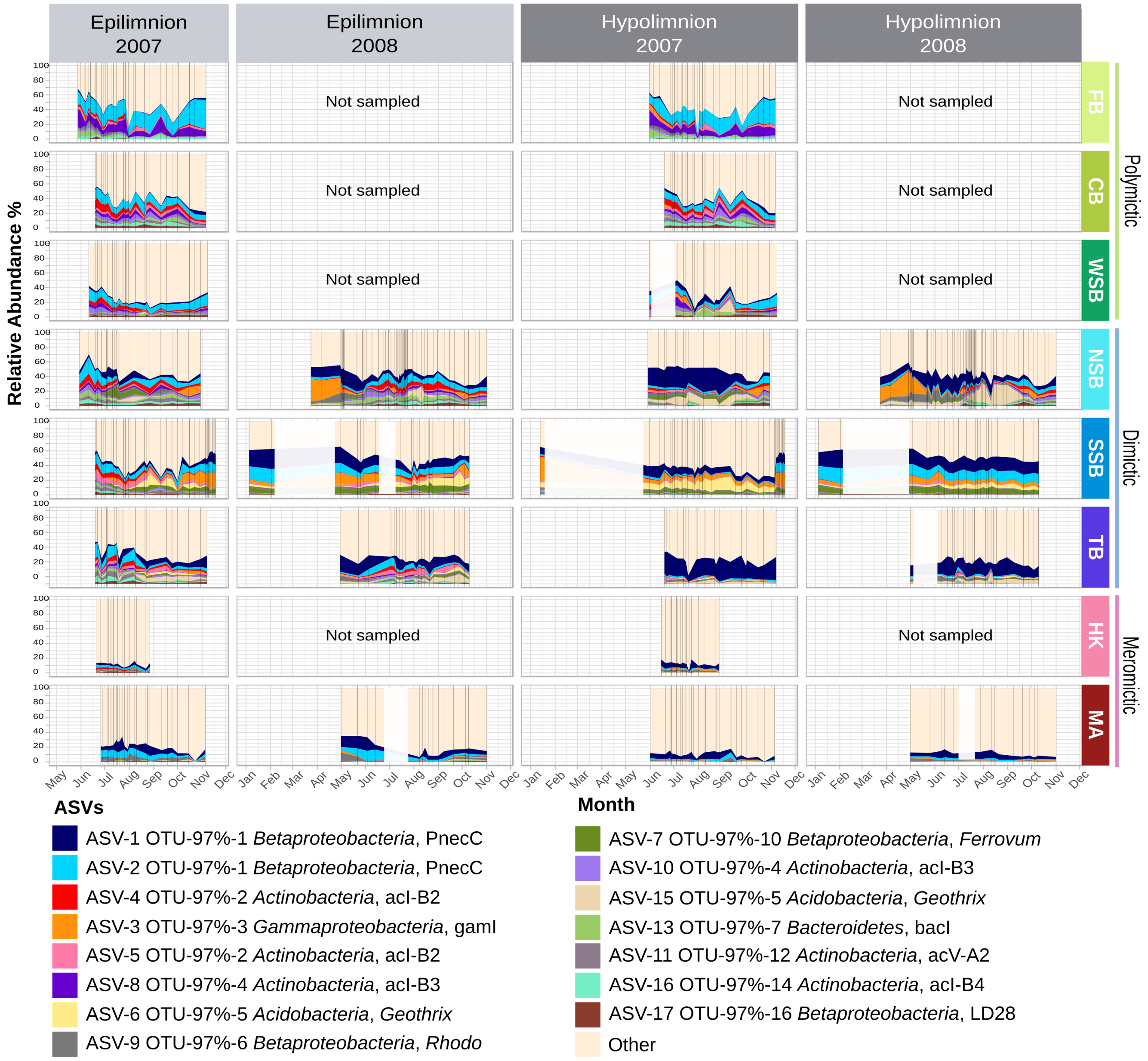


**Fig. S3. Abundance of the fifteen most abundant ASVs in the whole dataset.** Vertical lines indicate sampling points. We used linear interpolation between real samples. Periods interpolated over 30 days are masked by a white rectangle. The ASV 'Other' includes the sum of the abundances of the ASVs that were not plotted individually.

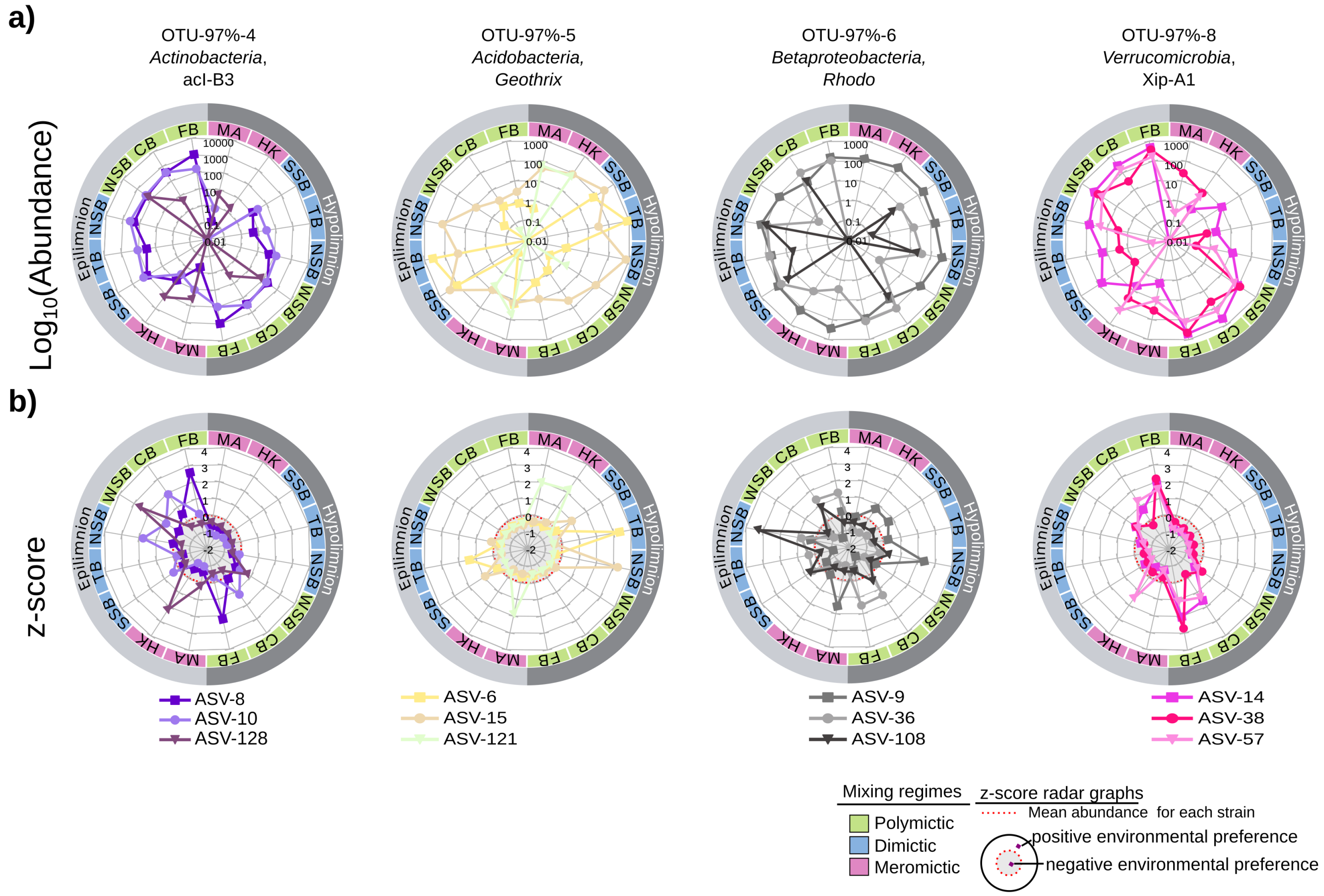


**Fig. S4. Environmental preferences for OTUs-97% numbers 4, 5, 6 and 8.** The radar plots show the abundance (logarithmic scale, a) and z-score (b) of their three most abundant ASVs. The abundances of the ASVs inform about their real distribution. The colors of the inner ring represent the mixing regime of the lake: green for polymictic, blue for dimictic and red for meromictic. The shades of grey in the outer ring indicate the lake layer (epilimnion and hypolimnion). Zero values have been plotted as $\log_{10}$ (0.01). Z-scores indicate the environmental preference of each ASV. Since the z-score normalization was made independently for each ASV, z-scores cannot be compared quantitatively among ASVs. Red dotted circumferences in z-score plots indicate the mean of their abundances, separating positive (white background) and negative (grey background) environmental preferences as schematically represented at the bottom right of the figure. ASV's points above the dashed red line exhibit a positive preference for that environment and vice-versa.

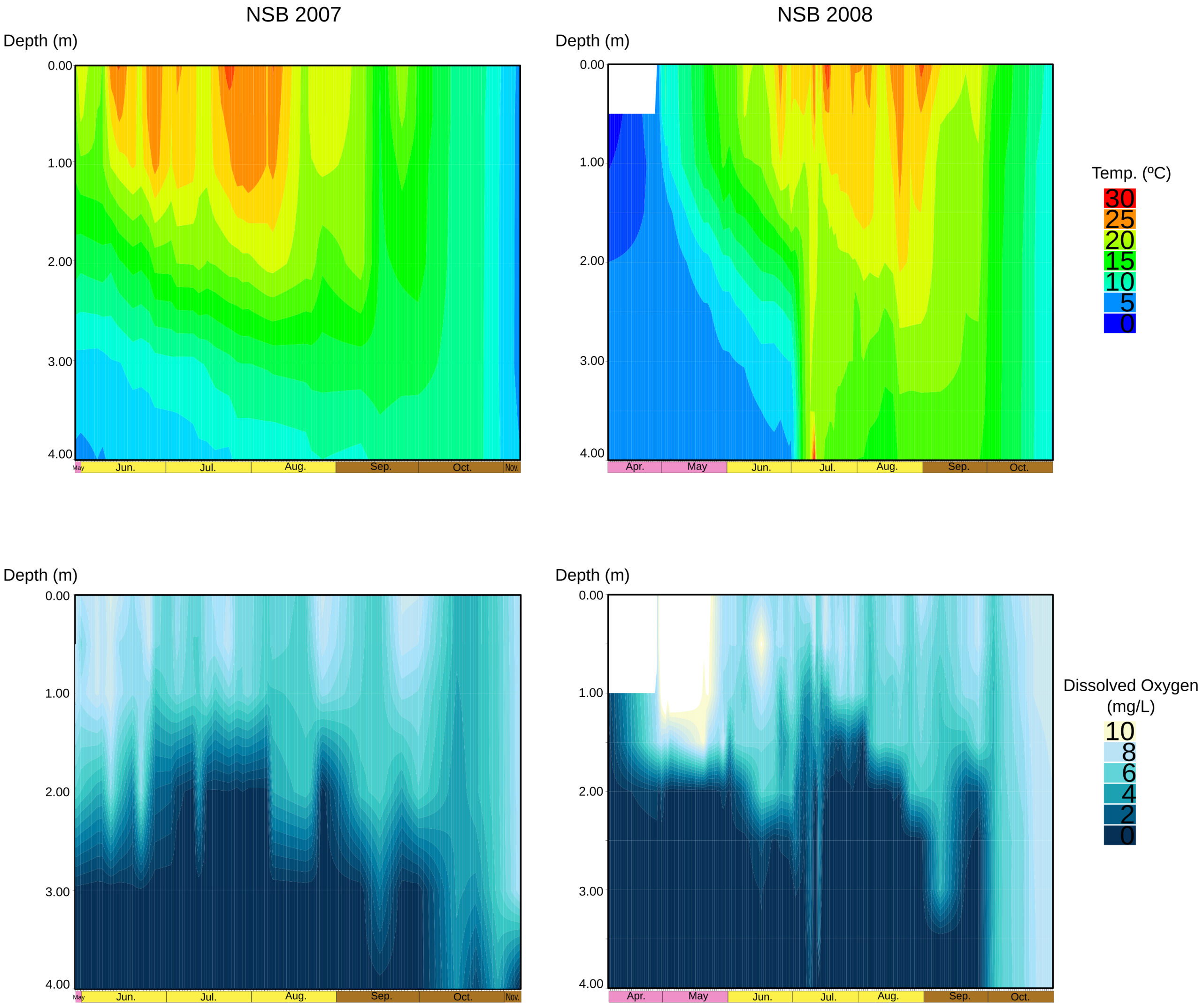


**Fig. S5. Temperature and dissolved oxygen profiles of NSB.** Homogeneous vertical temperature profiles correspond to mixing events. In 2008, there was an artificial mixing event from July 2 to July 10, when homogeneous temperature was observed. Black vertical lines in the different panels indicate sampling points.

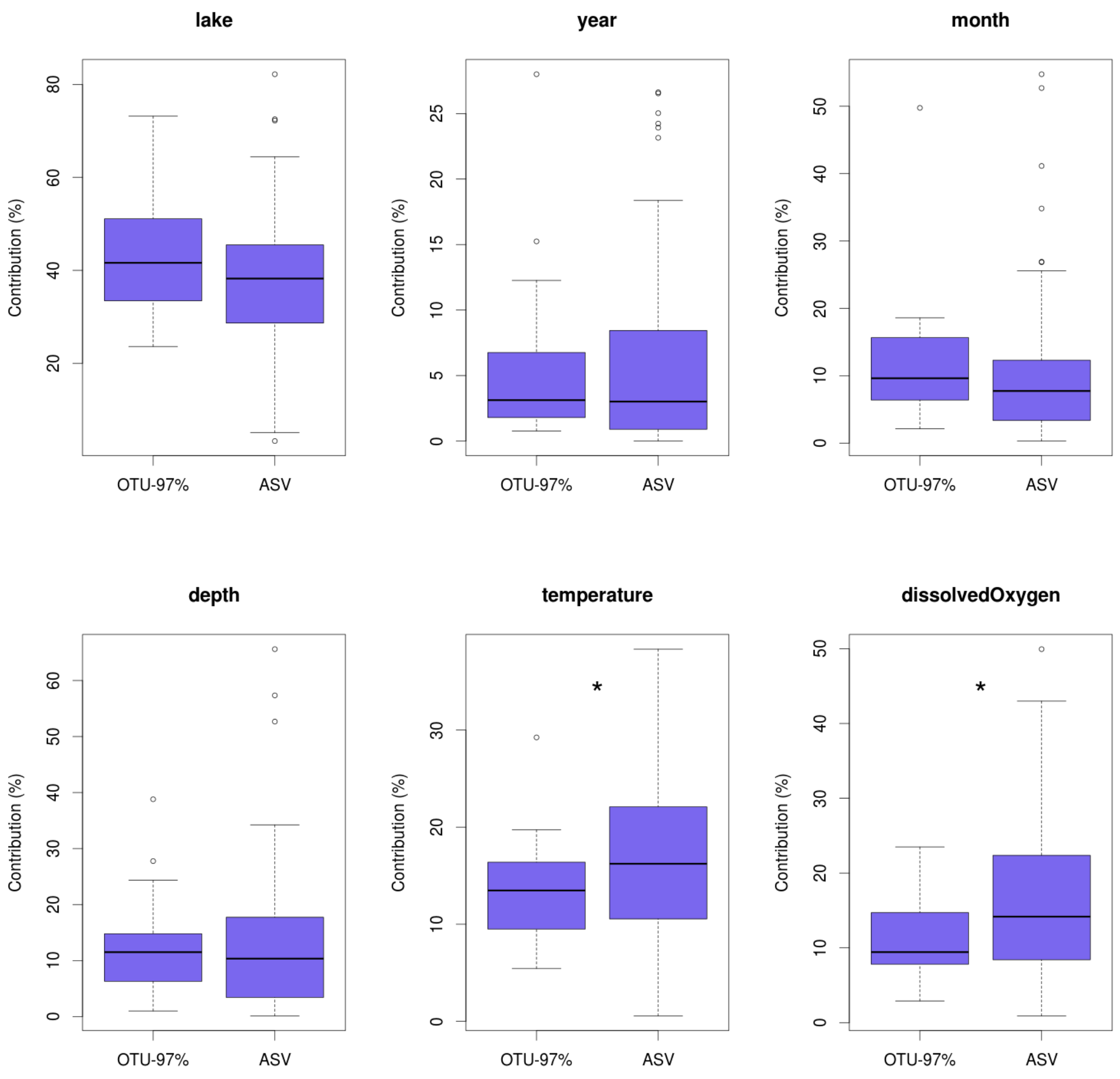


**Fig. S6. Influence of environmental factors on the distribution of the most stable OTUs-97% and their constituent ASVs.** Boosted regression trees were fitted to explain the abundance of OTUs-97% and ASVs as a function of six environmental variables. For each environmental variable, the boxplots show its influence (measured as its relative influence on reducing the loss function during model computation) in the distribution of both OTUs-97% and ASVs. Temperature and oxygen had higher (Welch's t test, $p = 0.041$ and $p = 0.014$, respectively) influences on the distribution of ASVs than on the distribution of their parent OTUs-97%.

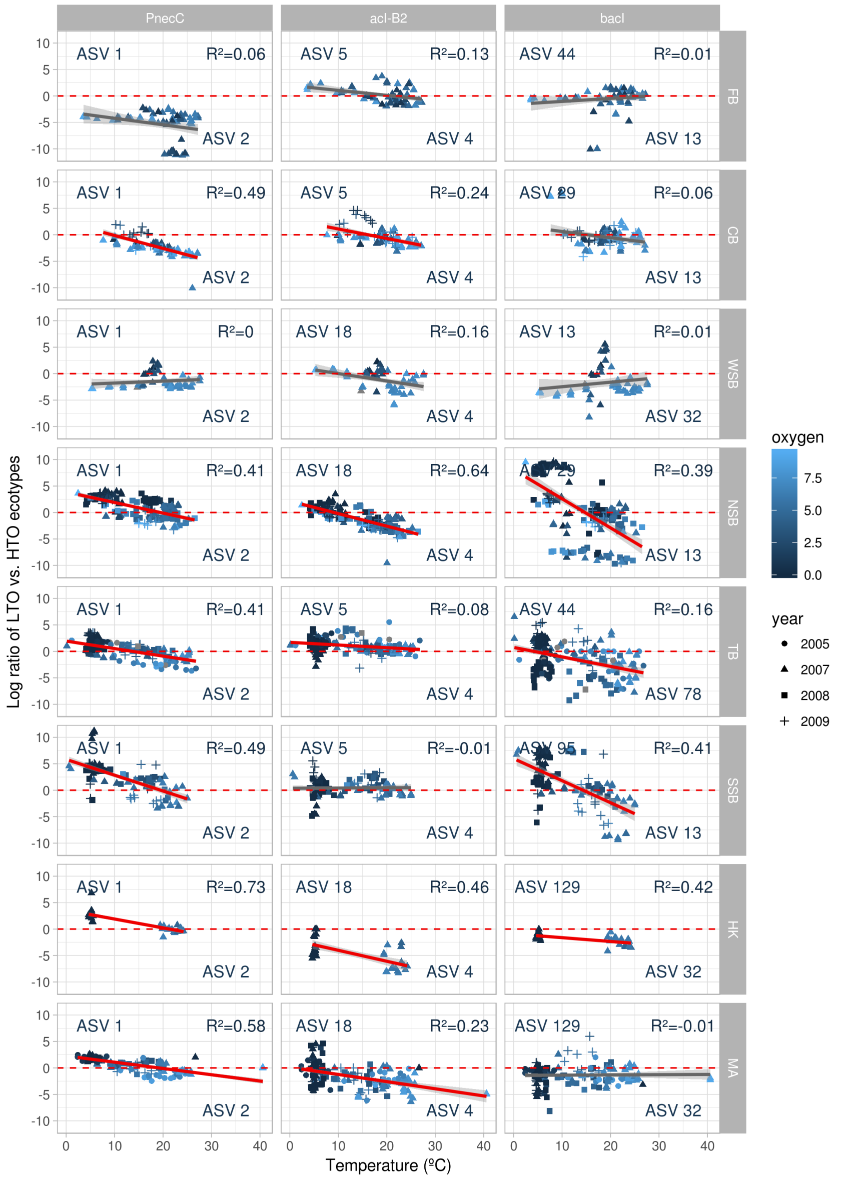


**Fig. S7. Relative abundance of low temperature/oxygen (LTO) versus high temperature/oxygen (HTO) ecotypes of the model community in samples from all the lakes, plotted as a function of water temperature.** Symbol color indicates dissolved oxygen concentration in shades of blue. Symbol shape indicates sampling year. Samples above the dashed red line are dominated by LTO ecotypes, and vice-versa. Significant linear regressions (FDR < 0.05) and 95% confidence intervals are shown with a red line and a shaded grey area, respectively. Non-significant linear regressions are shown with a grey line.

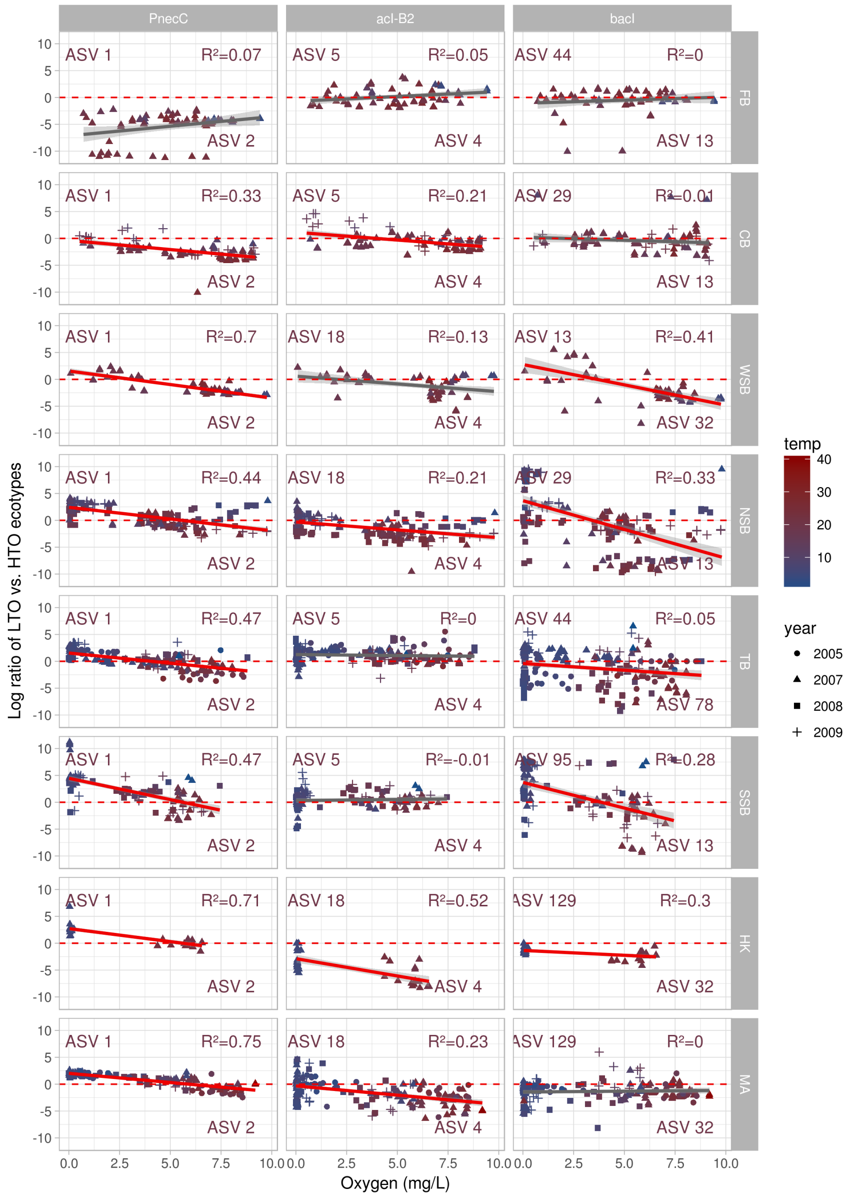


**Fig. S8. Relative abundance of low temperature/oxygen (LTO) versus high temperature/oxygen (HTO) ecotypes of the model community in samples from all the lakes, plotted as a function of dissolved oxygen concentrations.** Symbol color indicates water temperature from blue (low temperature) to red (high temperature). Symbol shape indicates sampling year. Samples above the dashed red line are dominated by LTO ecotypes, and vice-versa. Significant linear regressions (FDR < 0.05) and 95% confidence intervals are shown with a red line and a shaded grey area, respectively. Non-significant linear regressions are shown with a grey line.
